# The manipulation of spinal motor neuron axonal growth promotes spinal cord repair

**DOI:** 10.1101/2021.11.13.468456

**Authors:** Feng Wang, Xin-Ya Fu, Mei-Mei Li, Xing-Ran Wang, Ji-Le Xie, Jin-Jin Ma, Yan-Xia Ma, Saijilafu

**Affiliations:** Orthopaedic Institute, Medical College, Soochow University, Suzhou, Jiangsu, China; Department of Orthopaedic Surgery, the First Affiliated Hospital, Soochow University, Suzhou, China

**Keywords:** spinal motor neurons, axon growth, spinal cord injury, Pten, functional recovery

## Abstract

The loss of motor function in patients with spinal cord injury (SCI) is primarily due to the severing of the corticospinal tract (CST). Spinal motor neurons are located in the anterior horn of the spinal cord, and as the lower neurons of the CST, they control voluntary movement. Furthermore, its intrinsic axonal growth ability is significantly stronger than that of cerebral cortex pyramid neurons, which are the upper CST neurons. Therefore, we established an axonal regeneration model of spinal motor neurons to investigate the feasibility of repairing SCI by promoting axonal regeneration of spinal motor neurons. We demonstrated that conditionally knocking out *pten* in mature spinal motor neurons drastically enhanced axonal regeneration *in vivo*, and the regenerating axons of the spinal motor neurons re-established synapses with other cells in the damaged spinal cord. Thus, this strategy may serve as a novel and effective treatment method for SCI.

## Introduction

The loss of motor function in patients with spinal cord injury (SCI) is primarily due to the severing of the corticospinal tract (CST). The descending fiber of the CST is orientated from the pyramid neurons of the cerebral cortex and controls the voluntary movement of the skeletal muscle by innervating spinal motor neurons. However, because of their limited intrinsic axonal regeneration ability, spontaneous regeneration of CST axon fibers after SCI is difficult [1–3]. Therefore, over the past several decades, studies have largely focused on promoting CST axonal regeneration by enhancing the intrinsic growth ability of cerebral cortex pyramid neurons through the manipulation of the gene expression. For example, the deletion of several suppressor genes, such as *PTEN*, *SOCS3*, and *KLF4*, promotes axon regeneration in the injured CST [1–3]. However, even with this approach, the number and distance of regenerating axons are still insufficient to significantly improve the locomotive function of the damaged spinal cord. Thus, an alternative, more powerful, and more effective spinal cord repair strategy is imperative.

After SCI, spinal motor neurons, which reside below the lesion site, can still survive for a prolonged period and maintain their inherent synaptic connections and function. For example, after total spinal cord transection in rats, both the deep and superficial reflexes of the lower spinal cord are preserved. Moreover, in primates, including humans, the inferior deep reflex can be restored after SCI. As the lower motor neurons of the CST pathway, spinal motor neurons, together with the CST, control the voluntary movements of skeletal muscles. Additionally, the intrinsic regenerative ability of spinal motor neurons is significantly greater than that of its upper neurons, the cerebral cortex pyramid neurons. Therefore, if we can promote the axonal regeneration of spinal motor neurons to grow over the lesion area and re-establish synaptic connections with other downstream nerve cells in the spinal cord, motor functional circuits can be restored. If successful, it will help us to develop novel and effective treatments for SCI. On the other hand, spinal motor neurons extend their descending axons to innervate skeletal muscles in the periphery. Although motor neurons possess robust regenerative capabilities, because of the atrophic changes in chronically denervated Schwann cells in distal nerves and muscular atrophy, their functional recovery following peripheral nerve injury remains unsatisfactory in humans, especially with proximal lesions. Thus, promoting axonal regeneration of spinal motor neurons by manipulating gene expression may also serve as a powerful tool to genetically dissect motor axonal regeneration *in vivo*. Unfortunately, to the best of our knowledge, there have been few studies on the direct manipulation of spinal motor neuron intrinsic axonal growth ability to promote regeneration.

Here, we report a new therapeutic approach for spinal cord repair by promoting axonal growth of spinal motor neurons. Using Cre-Lox recombination techniques, we knocked out the *pten* (phosphatase and tensin homolog [PTEN]) gene in spinal motor neurons via a local Cre-expressing adeno-associated virus 2/9 (AAV2/9-Cre) injection into the anterior horns of tdTomato^f/−^/pten^f/f^ transgenic mice. Our data showed that conditionally knocking out *pten* in mature spinal motor neurons drastically enhanced their axonal regeneration in the injured spinal cord. Furthermore, we demonstrated that the regeneration of the axons of spinal motor neurons re-established synapses with other spinal cells. Using this approach, we also performed sciatic nerve injury after tdTomato-labeling motor axons and EGFP-labeling sensory axons. The regenerative axon length measurement provided solid evidence that the axon regeneration ability of motor neurons was significantly weaker than that of sensory neurons. Thus, our study not only established a powerful *in vivo* model to study spinal motor neuronal axon regeneration but also provided a novel strategy to repair SCI.

## Materials and methods

### Animals

All animals were handled following the animal protocol approved by the Institutional Animal Care and Use Committee of Soochow University. The Rosa-tdTomato reporter mice were kindly provided by Prof. Fengquan Zhou Laboratory of Johns Hopkins University, and the PTEN^f/f^ mice were a generous gift from the Prof. Yaobo Liu Lab of Soochow University. We mated the Rosa-tdTomato^f/−^ reporter mice with the PTEN^f/f^ mice and obtained the Rosa-tdTomato^f/−^/PTEN^f/f^ transgenic mice. The AAV2/9-Cre (1.0 × 10^12^ genomic copies per ml) was purchased from Hanbio (Shanghai, China).

### AAV2/9-Cre virus injection

To trace the spinal motor neurons, Rosa-tdTomato^f^/^-^ reporter gene mice were deeply anesthetized using intraperitoneal injection of a mixture of ketamine (100 mg/kg) and xylazine (10 mg/kg), and a small laminectomy was performed at lumbar 1 (L1) to expose the underlying the intumescentia lumbalis. Then, a 1.0 μL AAV2/9-Cre virus was slowly microinjected into the anterior horn with the Picospritzer III, a pneumatic-controlled injection system. A single injection was made at the coordinates of 0.7 mm deep and 0.5 mm lateral to the dorsal median sulcus. After suturing the muscle and skin tissue, mice were placed on a 35.0°C heating pad until they were fully awake.

### Immunohistochemistry staining

Two weeks after the intraspinal virus injection, the spinal cord, dorsal root ganglion (DRG), sciatic nerve, and gastrocnemius muscle were collected and fixed with 4% PFA at 4°C overnight. The samples were then processed into frozen sections as previously described [3]. For immunohistochemistry staining, sections were blocked in 0.1 M PBS solution containing 10% FBS and 0.5% Triton X-100 for 2 h and incubated with primary antibodies overnight at 4°C. After washing with PBS three times, sections were incubated with secondary antibodies, Alexa Fluor 568 (1:1000, Life Technologies) or Alexa Fluor 488 (1:1000, Life Technologies) for 1 h at room temperature. To quantify the number of axons of AAV2/9-Cre virus-infected motor neurons in the sciatic nerve, cross- and longitudinal sections of the sciatic nerve were stained with Tuj-1 (1:500, sigma, T5076) and ChAT (1:50, Millipore, AB144P). The numbers of Tuj-1-positive axons, ChAT-positive motor axons, and tdTomato-expressing axons were quantified. For visualization of the neuromuscular junction (NMJ), frozen longitudinal sections of the gastrocnemius muscles were stained as previously described [4], and the NMJ was defined by the degree of overlap between acetylcholine receptors (AChRs) stained with labeled α-bungarotoxin (α-BTX) and the axon terminals labeled by tdTomato-fluorescent protein.

### Spinal cord crush injury and axonal growth quantification

Under deep anesthesia, a small laminectomy was performed at L1 to clearly expose the spinal cord. The spinal column was fixed using toothed forceps, and the whole spinal cord was crushed with the modified 5^#^ fine forceps for 1 s. According to the method described above, the AAV2/9-Cre virus was injected into the spinal cord anterior horn at the caudal or rostral site of the crush injury. The bladder was squeezed and emptied every day until transcardially perfused. After 2 or 4 weeks of spinal cord crush injury, the mice were perfused transcardially with 4% PFA in 0.1 M PBS. The spinal cord was dissected out, and the cryostat was used to cut 12-μm sagittal sections. Then, the number of tdTomato-labeled axon fibers at a specified caudal or rostral distance from the lesion area was calculated according to the areas of the spinal motor neurons that were labeled with tdTomato, and the evaluated slices were averaged. Additionally, the vGlut-1 (1:200, SySy, Germany) antibody was used to visualize the pre-synaptic structure in the sagittal section of the spinal cord.

### Electroporation of adult mouse DRG neurons

As described in our previous studies [6, 7], the DRG of the L4 and L5 were exposed using laminectomy. Then, 1.0 μl EGFP plasmids (2.0 ug/ml) were microinjected into the DRG of the L4 and L5 using a glass needle. DRG electroporation was performed using a tweezers-like electrode and ECM830 (five 15-ms pulses at 35 V, with a 950-ms interval) immediately after injection. Subsequently, the wound was closed, and the mice were placed on a 35°C heating pad for recovery.

### Sciatic nerve injury and axonal length quantifications

Four days after DRG electroporation and intraspinal virus injection, the mice were anesthetized again and the sciatic nerves crush injury was performed as described previously [5, 6]. A further 3 days later, mice were killed by cardiac perfusion of 4% PFA in 0.1 M PBS (pH 7.4) and the whole sciatic nerve segment was carefully dissected to fix overnight in 4% PFA at 4.0°C. The distance from the crush injury site to the distal axon tip was measured in all EGFP- and tdTomato-labeled nerve fibers.

### Behavioral analysis

To analyze the functional recovery of the spinal cord, the Basso mouse scale (BMS) open-field locomotion rating score was used from days 1 to 28 as previously described [7]. Before the analysis, the bladder was squeezed and emptied, and the animals were placed under test conditions and allowed to move freely for 30 min to adapt to the environment. BMS scores were independently evaluated by an experimenter who was blinded to group allocation.

### Statistical analysis

Data are presented as means ± standard error of the mean (SEM), using Graphpad Prism 5. Axon number and length were quantified and t-tests were used to compare between the two groups. \**p* < 0.05, \*\**p* < 0.01, \*\*\**p* < 0.001, or non-significant difference (n.s.) denoted the significance thresholds.

## Results

### AAV2/9-Cre virus injection successfully infected spinal motor neurons

To label spinal motor neurons, we intraspinally microinjected the AAV2/9-Cre virus into the anterior horn of Rosa-tdTomato reporter mice (Figure 1A). One week later, we observed many cells in the sagittal and transverse sections of the spinal cord that expressed tdTomato red fluorescent proteins around the injection site (Figure 1B and C). To confirm whether these cells were spinal motor neurons, we performed immunofluorescence staining with SMN (1:500, Novus, NB100-1936) and ChAT (1:50, Millipore, AB144P), which are both specific markers of spinal motor neurons [8–12]. We found that many tdTomato protein-labeled cells located in the anterior horn also expressed SMN and ChAT proteins (Figure 1D–G). Thus, our data indicated that the locally injected AAV2/9-Cre virus successfully infected spinal motor neurons. We then further traced their axons, which extended into the sciatic nerve from the anterior horn, and calculated the percentage of tdTomato-positive axons. We stained the cross-sections of the sciatic nerve with Tuj-1 (1:500, Sigma, T5076) for all axons and ChAT antibody for motor axons. Results showed that tdTomato-labeled motor fibers were uniformly distributed in the sciatic nerve (Figure 2A and B), and the number of tdTomato-positive axons accounted for 25.63% ± 2.42% (n = 3) of the total number of Tuj-1-positive axons in the sciatic nerve (Figure 2C and D). In addition, the proportion of tdTomato-positive axons accounted for 72.6% ± 2.91% (n = 3) of the total number of ChAT-positive motor axons in the sciatic nerve (Figure 2E and F).

**Figure 1.**
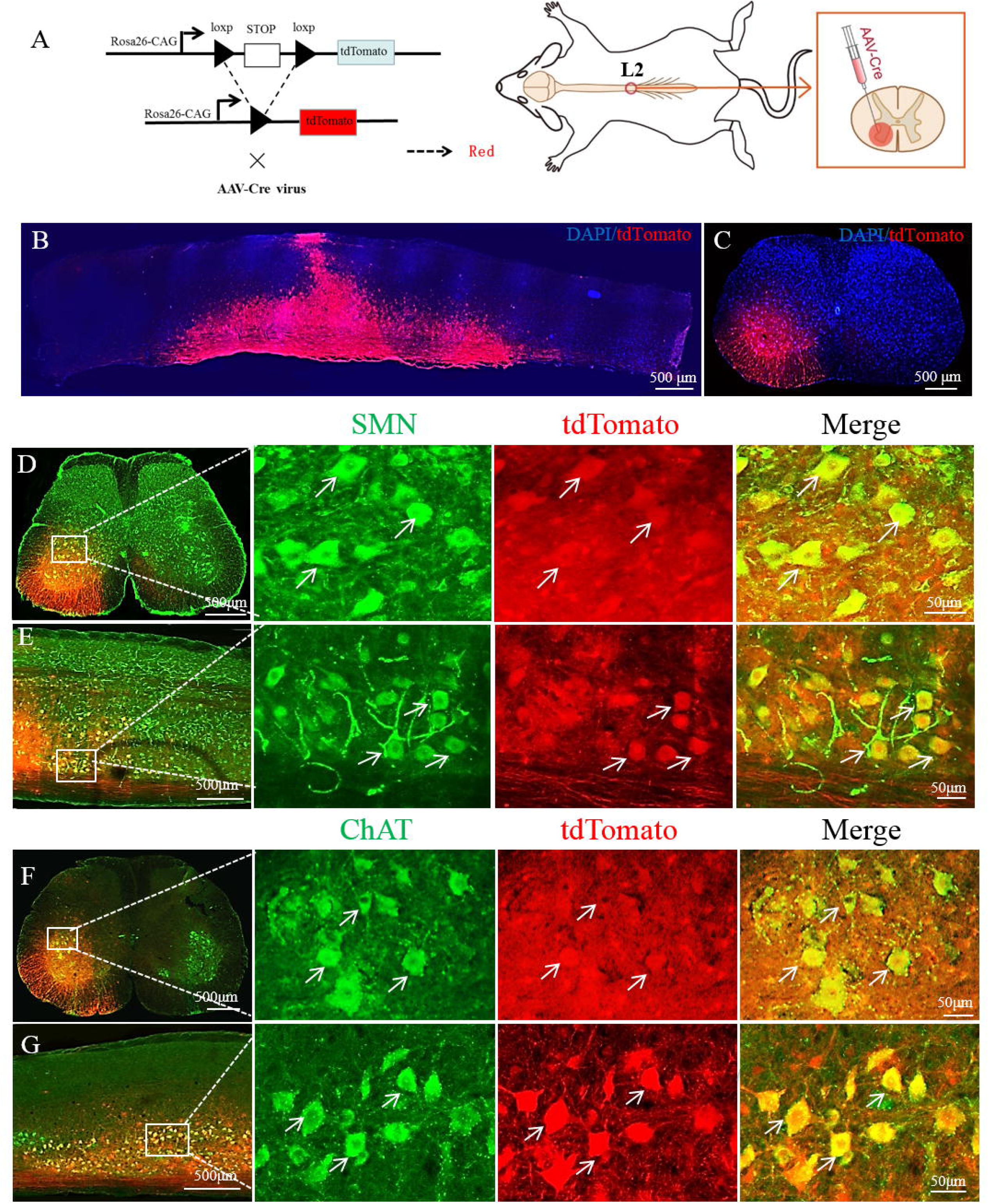
AAV2/9-Cre virus successfully infected spinal motor neurons. (A) Diagram showing injection of the AAV2/9-Cre virus into the spinal cord and Cre-induced expression of the tdTomato fluorescent protein in a Rosa-tdTomato reporter mouse. (B, C) Representative images of spinal cord sagittal sections showing that cells located in the anterior horn were successfully infected and expressed tdTomato fluorescent protein 2 weeks after AAV2/9-Cre virus injection. DAPI (blue) was used to label the nucleus and outline the spinal cord. (D–G) Immunofluorescence staining images showing that the tdTomato-positive cell also expressed SMN (D, E) and ChAT (F, G) proteins. Scale bar represents 500 μm (B–G) and 50 μm (squares in D–G).

**Figure 2.**
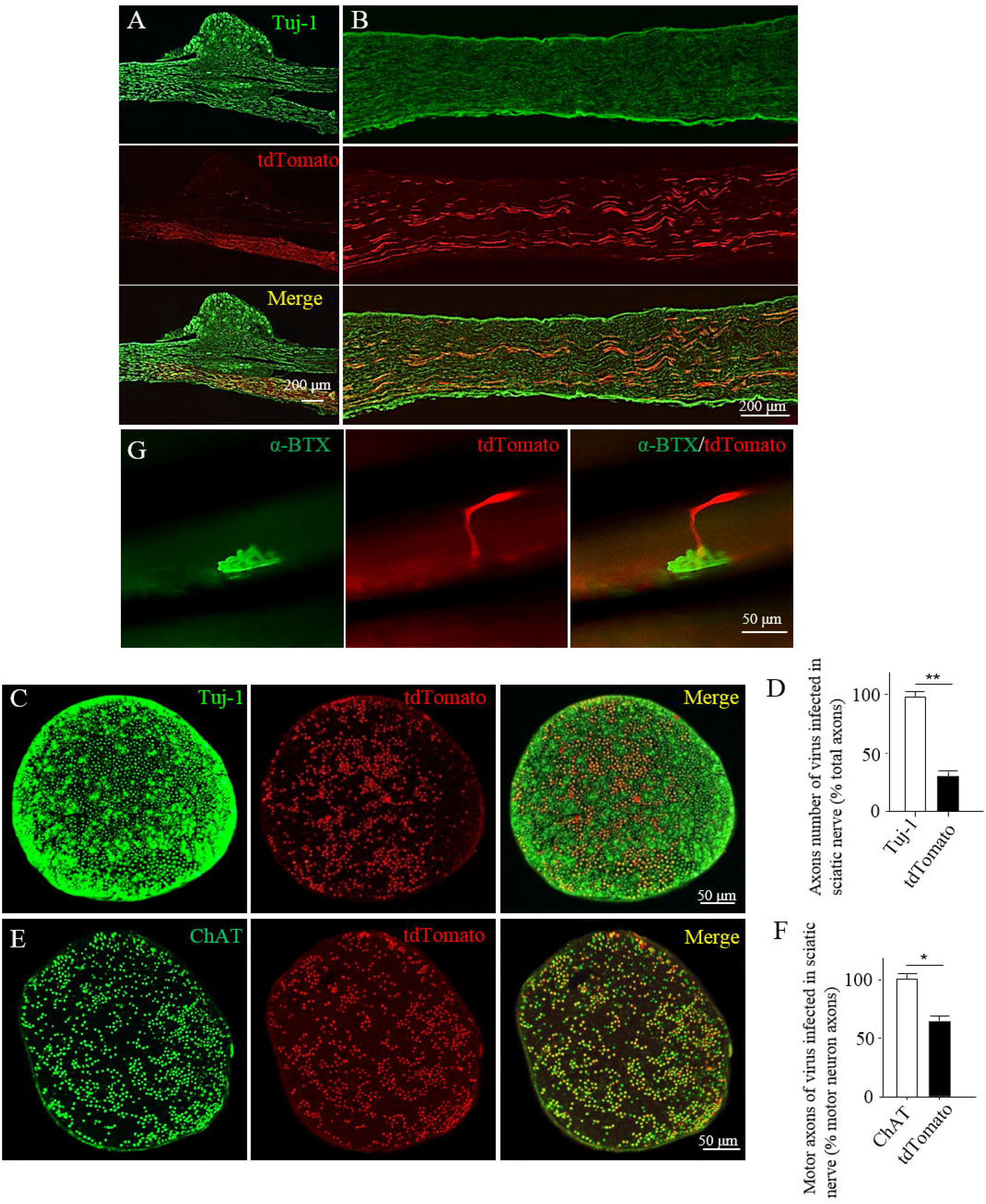
The whole length of spinal motor neurons was labeled with tdTomato fluorescent proteins. (A, B) Representative images from DRG and sciatic nerve longitudinal sections showing that the tdTomato protein-labeled axons (red) were Tuj-1 positive (green). (C, D) Quantification of axon number, including ChAT-positive and tdTomato-positive cells in the transection of the sciatic nerve. The rate of tdTomato-positive axons in ChAT-positive cells was calculated. Data are presented as means ± SEM. \*\**p* < 0.01,\**p* < 0.05. (G) Image of the gastrocnemius section showing tdTomato-positive motor axons forming the neuromuscular junction, together with the α-BTX-labeled AChR (green). Scale bar represents 50 μm.

The motor endplate is formed by the axon terminals of spinal motor neurons and AChRs on skeletal muscle fibers [13]. Nerve signals are transmitted to skeletal muscles through motor endplates and control skeletal muscle contractions. We stained the motor endplates with α-BTX (1:1000, Invitrogen, B13422), which is a specific marker of nicotinic AChRs [5, 14], in frozen sections of the gastrocnemius muscles (50 μm). We observed that tdTomato-positive axons terminals co-localized with α-BTX-labeled AChRs on muscle fibers to form the motor endplates (Figure 2G). Thus, we successfully and efficiently labeled intact spinal motor neurons from their soma to the motor endplate by locally injecting the AAV2/9-Cre virus into the anterior horn of the spinal cord.

### Suppression of PTEN expression in spinal motor neurons enhances axonal regeneration *in vivo*

Recently, it has been reported that the specific knockout of the *pten* gene markedly enhances the intrinsic axon growth ability of adult central nervous system (CNS) neurons [15, 16]. Therefore, to explore the feasibility of repairing SCI by promoting the axonal growth of spinal motor neurons, we knocked out the *pten* gene in spinal motor neurons using Cre-Loxp techniques. After the AAV2/9-Cre virus was injected into the anterior horn of Rosa-tdTomato^f/−^/PTEN^f/f^ transgenic mice, whole spinal cord crush injury was performed at the rostral or caudal site of the injection area. Four weeks later, we investigated tdTomato-labeled axons in the frozen spinal cord sections. Our results showed that compared with Rosa-tdTomato^f/−^ mice (Figure 3A), the Rosa-tdTomato^f/−^/PTEN^f/f^ group showed numerous tdTomato-labeled axonal growth over the lesion area (Figure 3B–E). Thus, these results suggested that knockout of PTEN significantly enhances the axonal regeneration ability of the spinal motor neuron and that repairing SCI via the manipulation of gene expression in spinal motor neurons is possible.

**Figure 3.**
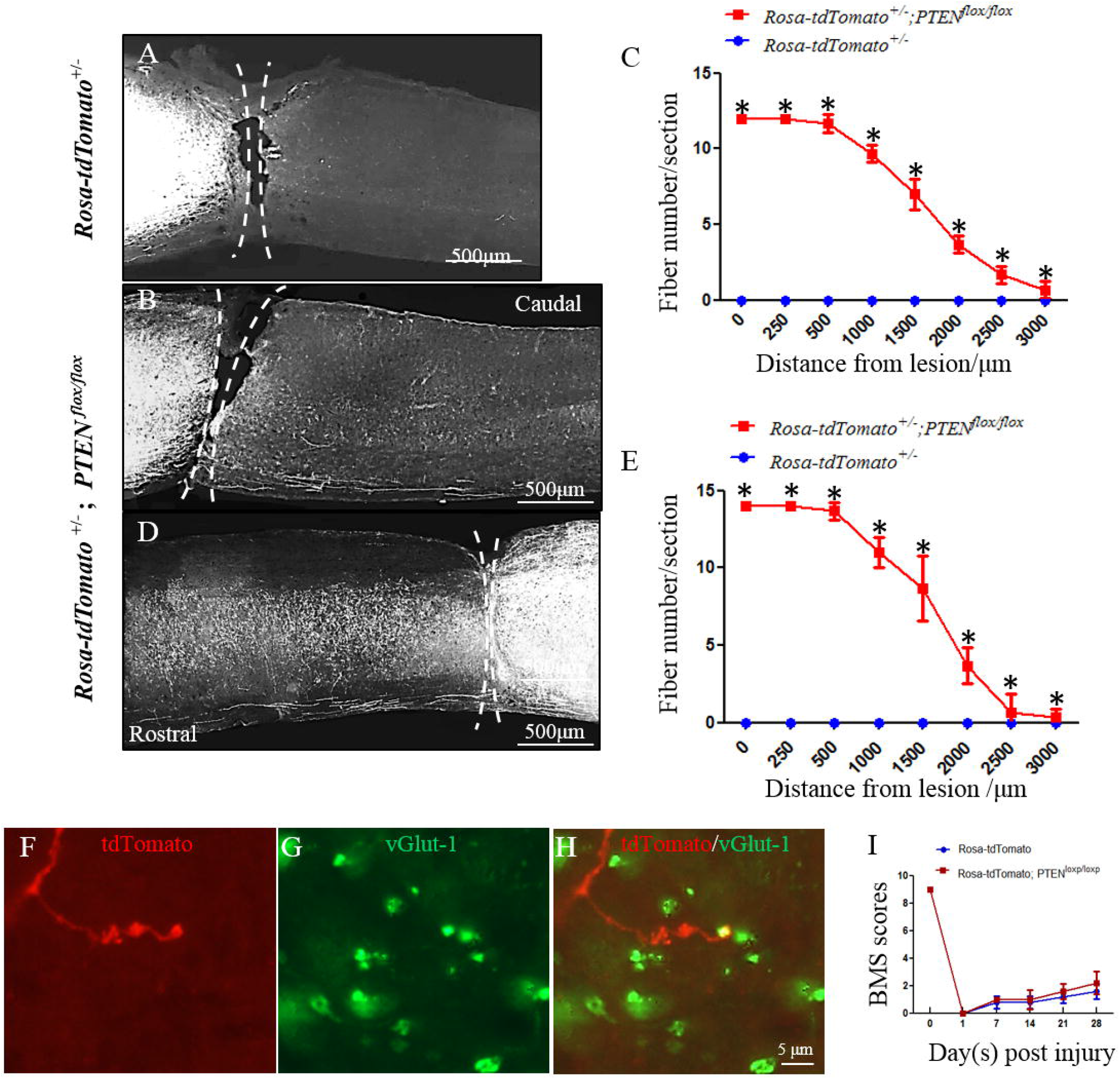
PTEN deletion in spinal motor neurons promoted axonal growth in the injured spinal cord. (A) AAV2/9-Cre virus was injected into the anterior horn of Rosa-tdTomato reporter mice, and a complete crush injury was performed. Four weeks later, there was no axonal growth over the lesion site. (B, C) PTEN deletion in the rostral spinal motor neurons of the spinal cord crush injury site promoted axonal growth over the lesion area to the caudal spinal cord. (C) is the quantification of (B). (D, E) PTEN deletion in caudal spinal motor neurons of the crush injury site promoted axonal growth over the lesion area to the rostral spinal cord. E is the quantification of (D). \**p* < 0.05. Scale bar represents 500 μm. (F–H) Representative images of sagittal spinal cord sections were stained with vGlut-1, and tdTomato-labeled axon ends overlapped with vGlut-1 buttons. (I) Quantification of BMS scores in Rosa-tdTomato^f/−^ and Rosa-tdTomato^f/−^/PTEN^flox/flox^ mice after SCI. Data are presented as means ± SEM for the two-group experiment. The experiment included five mice per group. Scale bars represent 5 μm.

Signaling information is transmitted from the upper neurons to downstream lower cells by synapses. Thus, we analyzed whether *pten* deletion-induced regenerating axons form synapses with other cells. Spinal cord sections were stained with vGlut-1, which is a presynaptic marker for excitatory synapses [11, 17, 18]. We found that tdTomato-labeled regenerating axon terminals overlapped with vGlut1-positive buttons (Figure 3F–G). Results showed that synaptic chemical molecules were accumulated and dispersed at the terminal of the regenerating axons, and the regenerated axons restored functional circuits. However, although there was no significant difference, *pten* depletion in spinal motor neurons showed higher BMS scores than those of controls at 21 and 28 days post-injury (Figure 3I). These results indicate that it is possible to manipulate gene expression in spinal motor neurons to promote axonal growth in the injured spinal cord.

### Axonal regeneration ability of motor neurons is weaker than that of sensory neurons

Our results showed that we successfully established an axonal regeneration model of spinal motor neurons. Therefore, we further investigated the axonal regeneration ability of spinal motor neurons themselves. We electroporated EGFP plasmid into the left L4/L5 DRGs of Rosa-tdTomato^f/−^ reporter mice as described previously [6], and simultaneously, we intraspinally injected the AAV2/9-Cre virus into the anterior horn (Figure 4A). Four days later, crush injury was applied to the left sciatic nerve of mice using fine forceps as described in our previous studies [6]. On day 7, the whole sciatic nerve was collected, followed by 4% PFA perfusion. The average regenerating axon length of EGFP-labeled sensory and tdTomato-labeled motor neurons in the sciatic nerve was 2823.23 μm for the sensory axon and 1971.65 μm for the motor axon (Figure 4B–E), which suggested that the intrinsic axonal regenerative ability of mature spinal motor neurons is dramatically weaker than that of peripheral sensory neurons.

**Figure 4.**
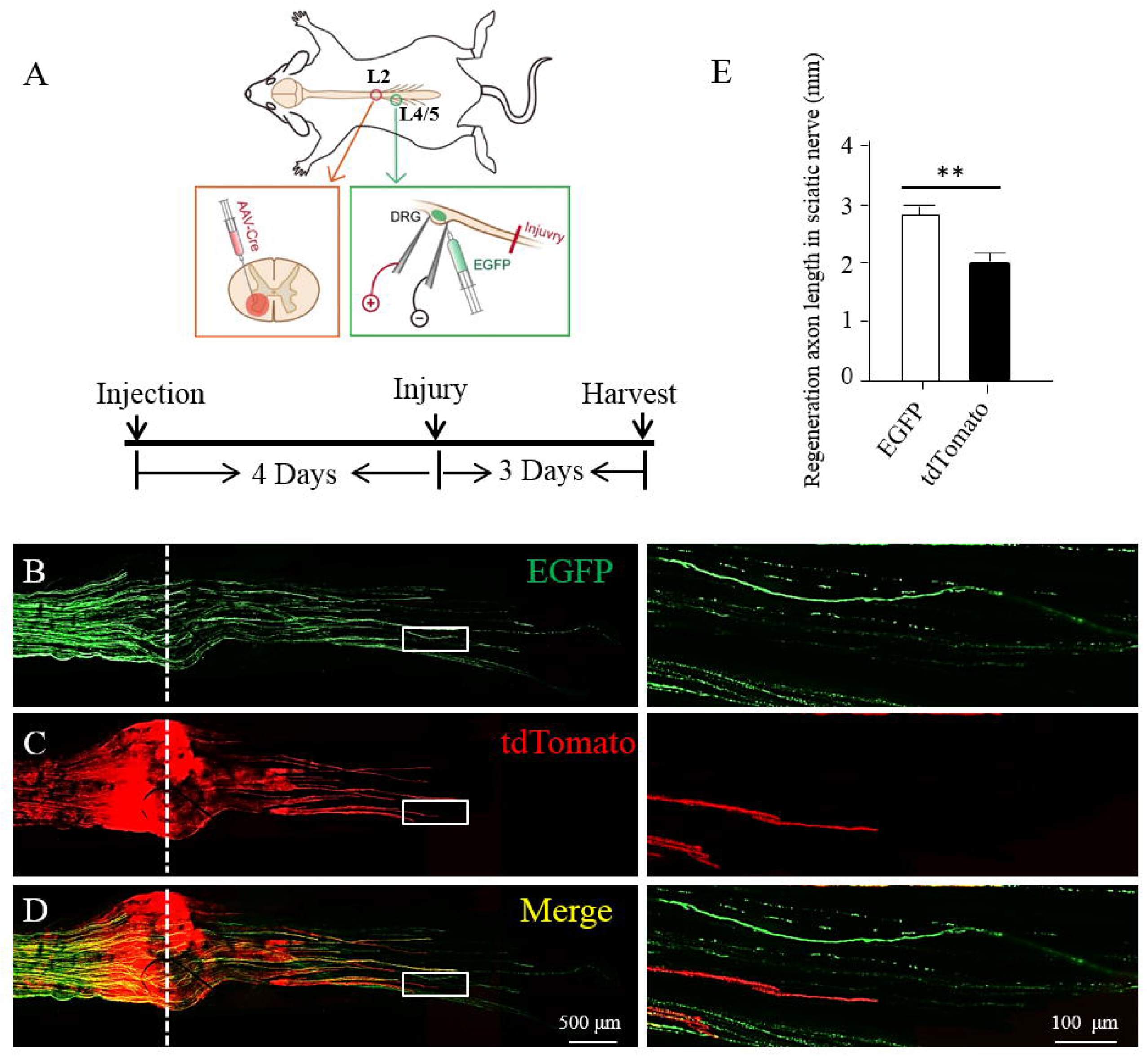
Axonal regenerative ability of spinal motor neurons was weaker than that of peripheral sensory neurons. (A) Schematic diagram of the intraspinal injection of the AAV2/9-Cre virus into Rosa-tdTomato reporter mice anterior horns and EGFP plasmid transfection into L4/5 DRGs by electroporation. (B–D) Representative images of the sciatic nerve section showing tdTomato-labeled motor neuron axons (red) and EGFP-labeled sensory axons (green). The regeneration length of motor axons was shorter than that of sensory neurons. The right panels are local magnifications of the graph on the left. The location of the crush injury site of the sciatic nerve is indicated by the dashed line. (E) Quantification of axon length in the sciatic nerve. The regeneration length of motor axons was shorter than that of sensory neurons. Data are presented as means ± SEM, using Graphpad Prism 5. \**p* < 0.05. Scale bar represents 500 μm (B–D) and 100 μm (B–D).

## Discussion

SCI affects millions of people worldwide and often causes permanent loss of motor, sensory, and autonomic function below the level of injury [1–3]. Furthermore, SCI involves high medical costs and economic burdens to families of patients and society in general. However, no effective treatment method for severe SCI has been made available to date. SCI often leads to axonotmesis, and subsequent regeneration of the injured axons becomes a key factor for functional recovery. Axonal regeneration re-establishes connections between neurons and downstream target cells and restores the integrity of neural circuits. The primary reason for axonal regeneration failure in the CNS is the diminished intrinsic axonal growth ability of mature CNS neurons and the inhibitory microenvironment, such as scar tissue and myelin sheaths. Although the CST controls voluntary movements, of the various nervous system tracts, it is the most difficult tract for natural axonal regeneration following injury [1–3]. As the lower motor neurons of the CST tract, spinal motor neurons play an essential role in voluntary movement because they represent the ultimate common pathway for delivering neural information to skeletal muscle fibers. All muscle movements, including breathing, walking, and fine motor skills, rely on spinal motor neuronal functions that transmit signals from the brain to muscle fibers. However, to the best of our knowledge, there have not been any reports or attempts to promote locomotor recovery by enhancing axonal regeneration of spinal motor neurons. Therefore, we established an axonal regeneration model of spinal motor neurons to investigate the feasibility of promoting axonal regeneration of spinal motor neurons to repair SCI. By microinjecting the AAV2/9-Cre virus into the L3–4 spinal cord ventral horn of Rosa-tdTomato^f/−^ reporter mice, we successfully labeled all spinal motor neurons with red fluorescence proteins. This indicated that it is possible to manipulate target gene expression in spinal motor neurons by intraspinally injecting the AAV virus. Moreover, we demonstrated that *in vivo* manipulation of gene expression in mature neurons with spatiotemporal control is an invaluable tool for studying motor neuron function. Additionally, our results showed that knockout of the *pten* gene in spinal motor neurons significantly promoted axonal growth over the injured area. The terminal of the tdTomato-labeled spinal motor axon also co-localized with vGlut-1-positive buttons, which suggested that the regenerating axons of spinal motor neurons re-established synaptic structure with other cells in the injured spinal cord. This finding indicated that it is possible to repair SCI by promoting axonal regeneration of spinal motor neurons.

Sciatic nerve injury is a well-established *in vivo* model of peripheral nerve regeneration [19–21]. The axons of spinal motor neurons enter the sciatic nerves via the ventral root and form a neuromuscular junction on muscle fibers. Here, we labeled motor axons with tdTomato fluorescence proteins by intraspinally injecting the AAV2/9-Cre virus into the L2 anterior horn of Rosa-tdTomato^f/−^ reporter mice. We also labeled sensory axons via electroporation of the EGFP plasmid into L4–L5 DRGs. Interestingly, we found that the regeneration length of the motor axon in the sciatic nerve was significantly shorter than that of the sensory axon. This indicated that the intrinsic axonal regenerative ability of the motor neuron was weaker than that of the sensory neuron. Using DRG neurons, numerous regeneration-associated genes (RAGs) that regulate the intrinsic axonal regeneration ability of sensory neurons were identified. To date, studies targeting intrinsic axonal growth ability (e.g., Pten, c-jun, ATF3, c-Myc, Lin28, LKB1) have produced highly promising results for peripheral sensory axonal regeneration [22–29]. However, despite extensive knowledge surrounding RAG expression and function in sensory neurons, few reports have demonstrated RAG-mediated axonal regeneration in spinal motor neurons, and thus, detailed molecular mechanisms by which these genes regulate motor nerve regeneration remain unclear owing to the lack of convenient and efficient animal models for genetic dissection of motor axonal regeneration *in vivo*.

Furthermore, direct damage to spinal motor neurons alone can lead to various motor dysfunctions, such as spinal muscular atrophy (SMA) and spinal nerve root avulsion. SMA is a disease characterized by the degeneration of spinal motor neurons, which leads to muscle weakness and muscular atrophy. Spinal nerve root avulsion is a severe nerve injury that predominantly occurs in work injuries, labor injuries, and traffic accidents, which result in the long-term loss of function of the affected limb. At present, there are no effective treatments for these conditions caused by the direct damage of spinal motor neurons. Moreover, the specific mechanism of axonal regeneration of motor neurons remains poorly understood. Therefore, it is of great significance to analyze and identify the regulatory roles played by relevant genes in the regeneration process of motor neurons. Furthermore, understanding the orderly regulation and intervention of injury repair is crucial.

In summary, using a newly developed spinal motor neuron regeneration approach, we demonstrated that SCI can be repaired by promoting spinal motor neuron axonal regeneration. Additionally, our results showed that compared with adult sensory neurons, spinal motor neurons possess weaker regenerative ability. Thus, our study not only established a powerful *in vivo* model to study spinal motor neuron axonal regeneration but also provides a novel strategy to repair SCI.

## Acknowledgement

This work was supported by the National Key Research and Development Program (No. 2020YFC1107402), the grant from the National Natural Science Foundation of China (No. 81772353 to Saijilafu), A Priority Academic Program Development of Jiangsu Higher Education Institutions, and Innovation and Entrepreneurship Program of Jiangsu Province.

## Author Contribution

F.W and S. designed the experiment. F.W performed the experiments and analyzed the data. F.W, XY. F, MM.L, XR.W, JL.X, JJ.M, YX.M and S. co-wrote the paper with all authors’ input.

